# Human nasal olfactory stem cell-derived extracellular vesicles improve the repair of rat nerves

**DOI:** 10.1101/2025.04.14.648811

**Authors:** Maxime Bonnet, Mostafa Seblani, Marie Witters, Tanguy Marqueste, Charlotte Jaloux, Philippe Morando, Patrick Decherchi, François Féron, Gaëlle Guiraudie-Capraz

**Author notes:** **Corresponding author:** Gaëlle GUIRAUDIE-CAPRAZ. These authors contributed equally to this work (Maxime Bonnet, Mostafa Seblani).

## Abstract

Damage to the peripheral nerve impairs quality of life. Despite the advances in surgery to facilitate nerve regeneration, complete recovery remains elusive. Human olfactory ecto-mesenchymal stem cells (OEMSC) have potential for the treatment of peripheral nerve injury through the release of extracellular vesicles (EV). The current research investigates the therapeutic effects of a venous bridge, filled with freshly purified or cryoconserved OEMSC-derived EVs after a peroneal nerve loss of substance.

Injured peroneal nerve was bridged with a vein into which freshly purified or cryoconserved EVs were injected or not. Nerve repair was analyzed by measuring locomotor function, muscle mechanical properties, muscle mass, axon number, and myelination. The EVs significantly increased locomotor recovery, maintained the contractile phenotype of the target muscle, and augmented the number of growing axons.

These results demonstrate that EVs display a positive effect on peripheral nerve regeneration, representing an alternative to cellular therapies for peripheral nerve repair.

## 1. Introduction

The peripheral nervous system (PNS) faces a myriad of challenges, with traumatic injuries constituting a substantial portion, accounting for 2.8% to 3% of all traumatic incidents ^1, 2^. The prevalence of these injuries is estimated to range between 13 and 23 cases per 100,000 people per year in developed countries ^3^. Such injuries often result in partial or total loss of sensory, motor, and autonomic functions within the affected body segments. Significant strides have been made in treating peripheral nerve injuries through advances in microsurgical procedures, including primary sutures, nerve grafts, and bioengineering innovations ^4^. Despite these technical achievements, the functional prognosis remains relatively poor especially in case of nerve defect, necessitating the exploration of complementary approaches.

Among these approaches, stem cell grafting has demonstrated effectiveness in regenerating injured peripheral nerves. Stem cells exert their regenerative influence through paracrine effects, involving the secretion of growth factors, the recruitment of glial cells and the establishment of a microenvironment conducive to nerve regeneration. While various stem cell types have been explored, challenges such as the difficulties of collection, the risks of teratoma formation and the inconsistent results for non-neural cells have spurred interest in a novel source: olfactory mucosa. The latter, which belongs to the PNS, is a site of permanent neurogenesis, fueled by progenitor cells and olfactory ecto-mesenchymal stem cells (OEMSC) ^5, 6^. These adult stem cells, members of the mesenchymal stem cell family, reside in the lamina propria and display strong mitogenic activity and a remarkable ability to differentiate into neural lineages. Notably, OEMSC are easily accessible in individuals of all ages and conditions, making them an attractive candidate for cell therapy experiments ^7^.

OEMSC have shown promise in various models, including paraplegia, Parkinson’s disease ^8^, deafness ^9^, and amnesia ^10^. The results of these different studies have shown: differentiation into glial cells and/or neural cells, upon transplantation into the nervous system ^8, 10, 11^, immunomodulatory actions ^12^ and the secretion of numerous factors involved in axon growth and guidance ^13^. These encouraging results have led us to evaluate the therapeutic potential of OEMSCs in PNS injury ^14, 15^. However, despite the recovery obtained in these studies, adult stem cell transplantation poses several problems such as the risk of neoplasia, the time required for culture and stem cell differentiation. Finally, stem cell differentiation is hardly required because somatic peripheral nerves do not contain interneurons and Schwann cells multiply and migrate without difficulty after trauma. Moreover, stem cells differentiation is not necessary for axonal regrowth, myelin formation, and functional recovery. Furthermore, the therapeutic advantages of stem cells primarily lie in the trophic and immune factors they secrete *via* their extracellular vesicles (EV) ^4, 16, 17^.

EV are formed by a lipid bilayer and contain numerous molecules (proteins, lipids, mRNAs, miRNAs, lncRNAs, DNA) that play a key messenger role in intercellular communication ^18^. They have physiological and pathophysiological roles in hemostasis, inflammation, transmission of information and biological molecules, cancer, metastasis or tissue regeneration ^19-23^. They also have many advantages: freezer storage, emergency availability, low immune response, reduced risk of embolism, no anarchic differentiation ^24-26^. Mesenchymal stem cell-derived EVs have been shown to exert beneficial effects on peripheral nerve regeneration by promoting (i) neo angiogenesis and the establishment of a microenvironment suitable to axonal nerve regeneration, (ii) neuro-regeneration and myelination, (iii) transport of growth factors, crucial for nerve regrowth (GDNF, BDNF, FGF-1, IGF-1, NGF) and their internalization by Schwann cells ^19^. Finally, EVs probably play an immunomodulatory role since their paracrine factors include more than 200 immunomodulatory proteins ^27^.

To further assess the therapeutic potential of OEMSC-derived EVs, we designed a study in which a 7 mm segment of rat peroneal nerve is removed, and a venous conduit is sutured to each nerve stump. Freshly purified and cryoconserved syngeneic OEMSC-derived EVs are inserted or not into the venous bridge, immediately after trauma. Transplanted and non-transplanted rats are compared to the Gold Standard animals in which the 7 mm nerve segment is immediately autografted in inverted position.

## 2. Materials and methods

The work has been reported in line with the ARRIVE guidelines 2.0.

### 2.1. Animals and ethical considerations

Experiments were performed on 35 adult male Sprague Dawley rats, weighing between 250-300g (Élevage JANVIER^®^, Centre d’Élevage Roger JANVIER, Le Genest Saint Isle, France), hosted two per cage in smooth-bottomed plastic cages in a laboratory animal house maintained on a 12:12-h light/dark photoperiod and at 22°C. Drinking water and rat chow (Safe^®^, Augy, France) were available *ad libitum*. Animals were housed during 2 weeks before the initiation of the experiment. All animals were weighed before each surgery and once a week until the end of the study.

Experiments were performed according to the French law (Decrees and orders N°2013-118 of 01 February 2013, JORF n°0032) on animal care guidelines and after approval by animal Care Committees of Aix-Marseille Université (AMU) and Centre National de la Recherche Scientifique (CNRS). The authorization number granted by the French Ministry of Higher Education, Research, and Innovation (MESRI) is APAFIS#41012. All persons were licensed to conduct live animal experiments and all room have a national authorization to accommodate animals (License n°B13.013.06). Furthermore, experiments were performed in accordance with the recommendations provided in the Guide for Care and Use of Laboratory Animals (U.S. Department of Health and Human Services, National Institutes of Health), with the directives 86/609/EEC and 010/63/EU of the European Parliament and of the Council of 24 November 1986 and of 22 September 2010, respectively, and with the ARRIVE (Animal Research: Reporting of *In Vivo* Experiments) guidelines.

After surgery, the animals were placed under a heat lamp until their thermoregulation was restored. The health status of the animals was monitored daily and animals showing signs of distress such as vocalization, lethargy, hyperactivity, significant weight loss (15 to 20%), and self-mutilation behavior were euthanized (no animal were excluded). They received subcutaneous injections (3 ml) of glucose-enriched saline solution to replace the fluid lost during the surgical procedure. Buprenorphine (0.03 mg/kg, 0.3 mg/ml, Bruprécare® Multi-dose, Axience Santé Animale SAS, Pantin, France) was administered subcutaneously before and after surgery daily for 3 days. A broad-spectrum antibiotic (Oxytetracycline, 400 mg/l, Sigma Aldrich, Saint-Quentin Fallavier, France) was diluted in the drinking water for 1 week to prevent any infections. Postoperative nursing care also included visual inspection for skin irritation or pressure ulcers, followed by cleaning of the hindquarters with soap and water and rapid towel drying of the fur.

### 2.2. Protocol design and experimental groups

After a two-week acclimation period, which involved familiarization sessions lasting one hour per day, three days per week, on - a walking corridor -, reference values (PRE-) were measured for Peroneal Functional Index (PFI) test which was used to follow the progress of recovery. Following the acclimation period, the animals were randomly assigned to five groups:

1. Control group (Control, n=7): No surgery was performed in this group.
2. Gold Standard group (GS, n=7): A 7-mm segment of the left peroneal nerve was excised and immediately autografted in inverted position.
3. Vein graft (VE) group (n=7): A 7-mm segment of the peroneal nerve was removed and a 1.5-cm vein conduit was immediately grafted between the two nerve stumps.
4. Vein graft and freshly isolated extracellular vesicles (VE-fEV) group (n=7): A 7-mm segment of the peroneal nerve was removed and a 1.5-cm vein conduit was immediately grafted between the two nerve stumps and filled with 5 million fresh EVs.
5. Vein graft and cryoconserved extracellular vesicles (VE-cEV) group (n=7): A 7-mm segment of the peroneal nerve was removed and a 1.5-cm vein conduit was immediately grafted between the two nerve stumps and filled with 5 million frozen EVs.

One week (W1) after surgery, animals underwent weekly assessments of sensory and motor recovery in the hindlimb. Evaluations were performed from Week 1 (W1) to Week 12 (W12) and compared to the baseline (PRE-) values. At the end of the twelve-week period the mechanical properties of the *Tibialis anterior* muscle were quantified using twitch measurements from a motor standpoint. Additionally, sensory aspects of the nerve were studied by analyzing type III-IV metabosensitive afferents in response to 1) electrically induced muscle fatigue (EIF) and 2) intra-arterial injection of potassium chloride (KCl) and lactic acid (LA) solutions. Then, animals were euthanized euthanized using an intraperitoneal overdose of pentobarbital sodium (1 ml) at a dosage of 280 mg/kg (i.p., Euthasol® Vet., Dechra Veterinary Products SAS, Suresnes, France) according to ethical recommendations.

Animals were randomly assigned to experimental groups using a randomized table in Excel. Each animal was identified by a number throughout the experiment to prevent disclosure of its experimental group. Group affiliations were revealed after data analysis conducted by blinded researchers. Potential confounders were minimized through standardization of treatment timing, housing conditions, and experimenter identity to ensure that all groups were similarly affected by potential variabilities.

### 2.3. Surgery procedure

Surgery of the peroneal nerve was performed as previously described ^15^. Animals were deeply anesthetized using 3% isoflurane (Isoflurin®, Axience Santé Animale SAS, Pantin, France) with a pre-anesthetic injection of buprenorphine (0.03 mg/kg, 0.85 mg/ml) administered 30 minutes prior. All surgical procedures were performed under aseptic conditions using binoculars.

The animals were positioned in a ventral position, and the left hindlimb was shaved and disinfected using Vétédine® solution 10% (Vetoquinol S.A., Magny-Vernois, France). In the GS, VE, VE-fEV, and VE-cEV groups, the peroneal nerve of the left limb was meticulously dissected from the surrounding tissues and cut to a length of 7 mm. In the GS group, the nerve segment was immediately replaced by the nerve segment in an inverted position and sutured (Ethilon 9-0, Ethicon) at the two free nerve stumps.

For the VE, VE-fEV, and VE-cEV groups, a 1.5 cm branch of the femoral vein was harvested from the contralateral side of the nerve injury. The harvested vein segment was washed in saline solution (NaCl 0.9%) and immediately used. The two nerve stumps were then inserted into the vein, leaving a 7 mm gap between the proximal and distal nerve stumps. To secure the graft, three or four 9-0 monofilament non-absorbable sutures (Ethilon® 9-0, Ethicon Inc., Johnson & Johnson, Somerville, New Jersey, USA) were used for each stump, along with biological thrombin/fibrinogen glue (Tisseel, Baxter^®^).

For the VE-fEV, and VE-cEV groups, 5 million fresh or stored OEMSC-derived EVs were suspended in 10 μl of phosphate-buffered saline (PBS). The EV were then injected into the vein using a 10 μl Hamilton syringe (Hamilton Company, Bonaduz, Switzerland).

### 2.4. Isolation and purification of extracellular vesicles from human olfactory ecto-mesenchymal stem cells (OEMSC-derived EV)

*Collection of olfactory mucosa biopsies*. OEMSCs were cultured based on a previously described protocol ^7, 28, 29^. Biopsies were unilaterally collected by an ENT surgeon, with informed consent from the patients participating in the Nose study. The biopsies were taken at the level of the middle turbinate arch using Morscupula forceps and an endoscope. Before the planned procedure and while the patient was under deep anesthesia, a 2 mm^2^ biopsy was obtained and transferred to a sterile tube filled with culture medium, penicillin, and gentamycin (alpha MEM Macopharma BC0110020).

*Culture of olfactory ecto-mesenchymal stem cells*. The olfactory mucosa, located on the nasal septum, was harvested from three healthy donors and placed in a Petri dish filled with DMEM/HAM F12. After triple washing to remove mucus, the biopsies were incubated in a Petri dish containing 1 ml of dispase II solution (2.4 IU/ml) for 1 hour at 37°C. The olfactory epithelium was then separated from the underlying lamina propria using a micro spatula. Once purified, the lamina propria was cut into small pieces using two 25-gauge needles and transferred to a 15-ml tube filled with 1 ml of collagenase NB5 (1U/ml, Nordmark Biochemicals). Following a 10-minute incubation at 37°C, the tissue was mechanically dissociated, and the enzymatic activity was halted by adding 9 ml of Ca- and Mg-free PBS. After centrifugation at 300g for 5 minutes, the cell pellet was resuspended in αMEM (Gibco Thermofisher©) supplemented with 10% platelet lysate (Macopharma®, Tourcoing, France) and seeded onto plastic culture dishes. The culture medium was refreshed every 2 to 3 days. Once reaching 80% confluence, typically between days 7 and 10, the cells were cryopreserved in liquid nitrogen for future use.

*Purification of OEMSC-derived EVs*. OEMSC were cultured in T175 flasks for amplification in a medium containing platelet lysate. On day 4, stem cells were washed with PBS to remove any contaminating vesicles present in the platelet lysate. Subsequently, the cells were cultured in αMEM (Gibco Thermofisher©) supplemented with insulin, transferrin, and selenium (ITS, 1%) for 4 additional days. Cell supernatants were collected and centrifuged at 300g for 5 minutes at room temperature and at 2,500g for 15 minutes at 20°C to obtain the vesicular supernatant. Extracellular vesicles were isolated by ultracentrifugation at 100,000g for 90 minutes at 4°C. Pellets of EVs were resuspended in PBS (less than 1 ml). EVs were purified using size exclusion chromatography columns (qEVoriginal Size Exclusion Column®, IZON, France). The purification procedure followed the instructions provided by the supplier. Purified OEMSC-derived EVs were either kept at 4°C for immediate use (referred as “freshly purified extracellular vesicles”, fEV) or cryoconserved at −80°C for 6 months (referred as “cryoconserved extracellular vesicles”, cEV).

### 2.5. Functional assessment of hind limb recovery

To evaluate functional changes, footprints were recorded and analyzed every week from W1 to W12 after the surgery, using a paper track method as previously described ^15^ and compared to PRE-values. Throughout assessments, experimenters involved in data collection were blinded to the treatment group, ensuring unbiased evaluations.

*PFI test*. The PFI was calculated using the formula described by ^30^ : PFI = 174.9 × [(ePL-nPL)/nPL] + 80.3 × [(eTS-nTS)/nTS] - 13.4. This index considers the parameters measured for both the normal (n) and operated (e) feet, including the footprint length (PL), which is the longitudinal distance between the tip of the longest toe and the heel, and the total toe spreading (TS), which is the cross-sectional distance between the first and fifth toes. The recovery rate of the PFI was assessed on a scale ranging from −100 to −13.4, where −13.4 represents normal function and −100 indicates total failure. Footprints were collected and analyzed from the first (W1) to the twelfth (W12) week post-surgery.

### 2.6. Electrophysiological recordings

At Week 12, rats were anesthetized using an intraperitoneal injection of Ketamine solution (75 mg/kg) and medetomidine (0.5 mg/kg). The following procedures and measurements were performed as previously described ^15^. In brief, the left peroneal nerve was dissected and carefully separated from the surrounding tissues over a length of 3-4 cm. A catheter was inserted into the right femoral artery and advanced to the bifurcation of the descending abdominal aorta. This catheter was used to transport chemicals (potassium chloride, KCl [20 mM in 0.5 ml of saline] and lactic acid, LA [1 mM in 0.1 ml of saline]) to the contralateral muscle. It also allowed free blood flow into the muscles of the lower left limb.

*Twitch measurement*. Two stimulation electrodes with an inter-electrode distance of 1 mm were placed on the surface of the peroneal nerve. The contractile response of the Tibialis anterior muscle to nerve stimulation was induced using a neurostimulator (S88K stimulator) that delivered rectangular single shocks (duration: 0.1 ms, frequency: 0.5 Hz). The response was measured using an isometric strain gauge (micromanometer 7001) attached to the tendon of the *Tibialis anterior* muscle. Several parameters, including amplitude (A), maximum contraction rate (MCR), and maximum relaxation rate (MRR), were recorded. MCR and MRR were normalized to the amplitude of the twitch (MCR/A and MRR/A, msec-1). The twitch data was recorded using the Biopac MP150 system (Biopac Systems Inc, Goleta, CA, USA), sampled at 2000 Hz, and analyzed using AcqKnowledge 3.7.3 software (Biopac Systems Inc.).

*Afferent activity*. A monopolar tungsten electrode was positioned under the *Tibialis anterior* nerve, which was immersed in paraffin oil. Nerve activity was recorded using a differential amplifier and filtered between 30 Hz and 10 kHz (P2MP®, 5104B, Marseille, France). The afferent discharge was recorded and analyzed using pulse window discriminators and the Biopac AcqKnowledge software. The response of muscle afferents was recorded after a 3-minute Electrically-induced muscle Fatigue (EIF) period and intra-arterial bolus injection of KCl (20 mM in 0.5 ml of saline) or LA (1 mM in 0.1 ml of saline) solutions. EIF was elicited using the S88K stimulator, which delivered pulse trains to the muscle surface electrode (pulse duration: 0.1 ms, frequency: 10 Hz, duty cycle: 500/1500 ms). The discharge rate of nerve afferents was averaged for a 30-second period before (regarded as baseline discharge) and after EIF or metabolite injection, and the increase in average afferent discharge rate was expressed as a percentage of the baseline discharge rate. A 20-minute recovery period was allowed between different interventions.

### 2.7. Euthanasia, muscular atrophy measurement and biopsy collection

At the end of the electrophysiological recordings, animals were euthanized using an intraperitoneal overdose of pentobarbital sodium (1 ml) at a dosage of 280 mg/kg (i.p., Euthasol® Vet., Dechra Veterinary Products SAS, Suresnes, France).

Following euthanasia, the left *Tibialis anterior* muscle was immediately harvested, weighed and preserved in a PBS solution containing 4% paraformaldehyde (PFA) then transferred in stored in sodium azide solution (0.1%) at 4°C for further histological investigation. The muscle weight was normalized by dividing it by the body weight, and the muscle/body weight *ratio* was calculated as an indicator of muscle atrophy.

Additionally, the peroneal nerves (n=4 per group) were excised, washed in phosphate-buffered saline (PBS) to remove contaminants, and stored in a PBS solution containing 4% paraformaldehyde until histological analysis. Some of the nerves (n=3) were previously immersed in a 2% glutaraldehyde solution containing PBS for 24 hours before being transferred to the 4% paraformaldehyde solution.

### 2.8. Immunohistochemistry, histology and quantification

*Immunohistochemstry*: For each group of animals, nerves were cut in three parts (proximal end, medial part, and distal end). Specimens fixed in PFA were processed in HistoGel^TM^ (Epredia^TM^ HG4000012, Thermo Fisher Scientific) then embedded in paraffin. After embedding, sections of 5 μm were performed using a vibrating blade microtome (Leica Biosystems, PRID: SCR_016495) and collected on coated slides. Sections were dewaxed in three changes of Histo-Choice® (Sigma Aldrich-Merck) for 10 min each, then rehydrated by transferring in decreasing concentrations of ethanol (100%, 95%, 70%, 50%), twice for 10 min each, and rinsed in two changes of distilled water, for 5 min each. The distal, medial, and proximal sections were immunostained with a mouse monoclonal antibody raised against the light chain of neurofilament protein (NF-L 70 KDa, Sigma Aldrich-Merck, dilution: 1:500). After washing, an appropriate biotinylated-conjugated secondary antibody was applied to the sections. The final staining step was performed using diaminobenzidine kit (DAB substrate kit, ab64238, abcam). Sections were delipidated in xylene and embedded in Eukitt mounting medium (Sigma Aldrich-Merck).

*Histology (p-Phenylenediamine staining)*: For myelin quantification, nerve specimens were washed three times for 5 min each in PBS and immersed into osmium tetroxide (2%) solution for 1 h. After 3 washes in PBS, 5 min each, samples were immersed in increasingly concentrated acetone solutions (50%, 35 min; 70%, 1 h; 95%, 1 h; 100%, 2 h, respectively). Samples were immersed in increasingly concentrated araldite solutions (50%, 3 h; 80%, 5 h, respectively). Specimens were immersed in araldite 100% and placed in heat chamber (80°C) during 12 h for resin polymerization. After inclusion, semithin sections (0.8 μm) were cut using an ultramicrotome and collected on coated slides. A solution of 1% p-phenylenediamine was prepared in absolute methanol and kept at room temperature in a closed dark glass jar protected from the light and evaporation 5 days before using it. After 5 days, the solution was filtered with a 0.2 μm nylon sterile filter (Millipore). Slides were stained by being immersed for 10 min in the staining solution, washed in tap water then in distilled water, and dried for 1h on a hot plate. Sections were mounted in Eukitt medium.

*Microscopy and quantification:* The stained sections were examined using an Apotome microscope (Apotome v2, AxioObserver Z1, Zeiss) associated with high-resolution camera (Camera CMOS Orca flash 4.0 v2, Zeiss, Oberkochen, Germany). The slides were digitized and analyzed with ImageJ (NIH) software. Axon numbers were counted for several sections in proximal, medial, and distal parts for each group of animals. Number of myelinated axons was assessed using a deep-learning tool by Axonet 2.0 (Goyal et al., 2023). The G-ratio (i.e., the ratio between the diameter of the axon and the outer diameter of the myelinated fiber) was calculated, using a semi-automated quantification using Myeltracer software ^31^ by assessing 100 fibers per section, randomly and blindly selected.

### 2.9. Statistical analysis

Results obtained from behavioral locomotor test (PFI), electrophysiological recordings (EIF, KCl and lactate injection), muscle properties (weight/body weight ratio, contraction properties) and histological data were compared between all experimental groups. Data processing was performed using Rstudio for statistical computing and graphics (Version 2024.04.2 Build 764, 2024). Significant differences were determined using non parametric tests since our data are not normally distributed ^32^. For the one-way experimental design, we applied the Kruskal-Wallis one-way analysis of variance ^33^, followed by Dunn’s test for post-hoc comparisons ^34^. For the two-way experimental design (experimental groups × time points), we utilized the Scheirer-Ray-Hare test, an extension of the Kruskal-Wallis test adapted for factorial designs ^35-37^ followed by Dunn’s test ^34^. Data were expressed as median ± SD. Difference was considered significant when p < 0.05.

## 3. Results

### 3.1. OEMSC-derived EVs improve locomotion but not muscle weight

After surgery, during which OEMSC-derived EVs were injected in the venous chamber, the locomotor behaviour of rats was measured and compared with the control and gold standard groups (**Figure 1A)**. The kinetics between the four groups are similar during the first 6 weeks post-surgery. The peroneal functional index significantly increased from W7 to W12 for rats receiving freshly purified EVs (fEV) compared with GS and VE animals. Statistically significant differences are observed at W7 (p<0.05), W8 (p<0.01), W9 (p<0.05), W10 (p<0.001), W11 (p<0.01), W12 (p<0.001) for GS *versus* VE-fEV and W7 (p<0.01), W8 (p<0.01), W9 (p<0.05), W10 (p<0.05), W11 (p<0.001), W12 (p<0.001) for VE *versus* VE-fEV. Peroneal functional index significantly increased for cEV-treated rats from W7, W12 (except at W10) when compared with Vein bridge-implanted animals (W7, p<0.05; W8, p<0.05; W9, p<0.05; W11, p<0.01; W12, p<0.01. When compared with GS rats, the cEV treatment demonstrates a significant improvement of the locomotion at W7 (p<0.05), W8 (p<0.05) and W12 (p<0.01). Three months after surgery, animals receiving extracellular vesicles, either freshly purified or cryoconserved, display a PFI value (−15.62 and −18.76, respectively) close to those of uninjured rats (−13.7). On average, the PFI values of the GS and VE rats - −39.86 and −41.18, respectively – remain far from control values. **Figure 1B** compares the *ratio* “muscle weight/body weight”. At Week 12, this ratio is significantly reduced for all four groups (p<0.001) when compared with the control value (dotted line). No difference is observed between the GS, VE, VE-fEV and VE-cEV groups.

**Figure 1.**
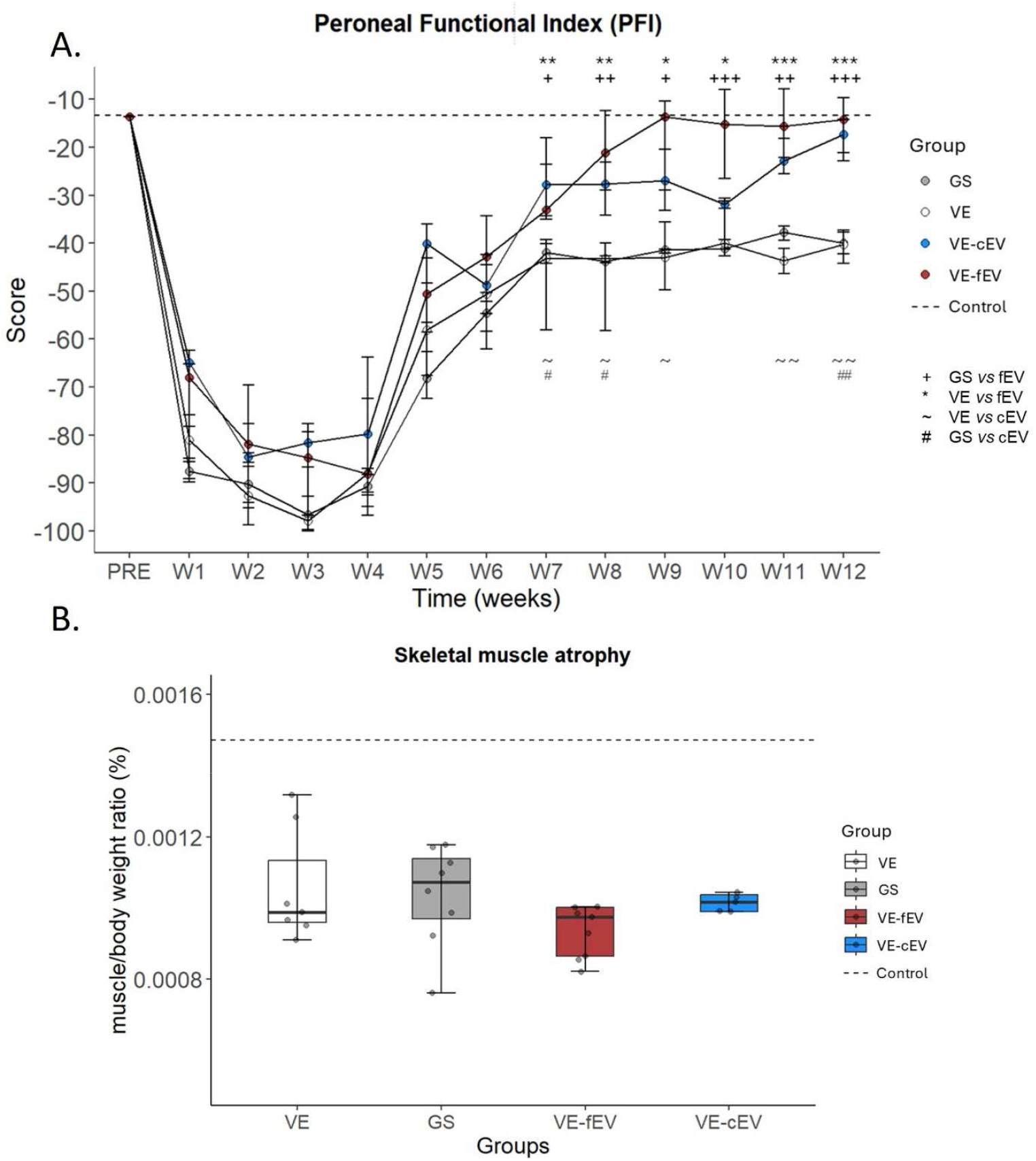
Peroneal Functional Index (PFI) and relative weight of the tibialis anterior target muscle. **(A)** The PFI was measured every week from W1 to W12 post-surgery. A significative improvement was observed from W7 to W12 in the fEV treated group when compared with VE and GS groups. A significative improvement is also observed in the cEV treated group at W7, 8, 11 and 12 when compared with GS, and at W7, 8, 9, 11 and 12 when compared with VE group. **(B)** The relative weight of the tibialis anterior was measured at W12 post-surgery. The muscle weight / body weight ratio was the same between the four groups. 1 symbol: p<0.05; 2 symbols: p<0.01; 3 symbols: p<0.001. Dotted line, mean value of Control. *GS, Gold Standard; VE, Vein; VE-cEV, Vein-cryoconserved extracellular vesicles; VE-fEV, Vein-freshly purified extracellular vesicles; W, Week*.

### 3.2. OEMSC-derived EVs partially modify the contractile phenotype of the target muscle

The peroneal nerve was electrically stimulated to elicit the contractions of the target muscle *Tibialis anterior*. The MCR/A *ratio* indicates that the phenotype of the target muscle is modified in VE and GS groups with a shift to a slower phenotype. Conversely, injected OEMS-derived EVs allow to maintain a phenotype close to a control situation (dotted line), particularly for the VE-fEV group, which significantly differs from the GS group (p<0.01) (**Figure 2A**). In addition, the fEV-treated animals display a significant improvement (p<0.05) of the MRR/A *ratio* compared to GS rats (**Figure 2B**).

**Figure 2.**
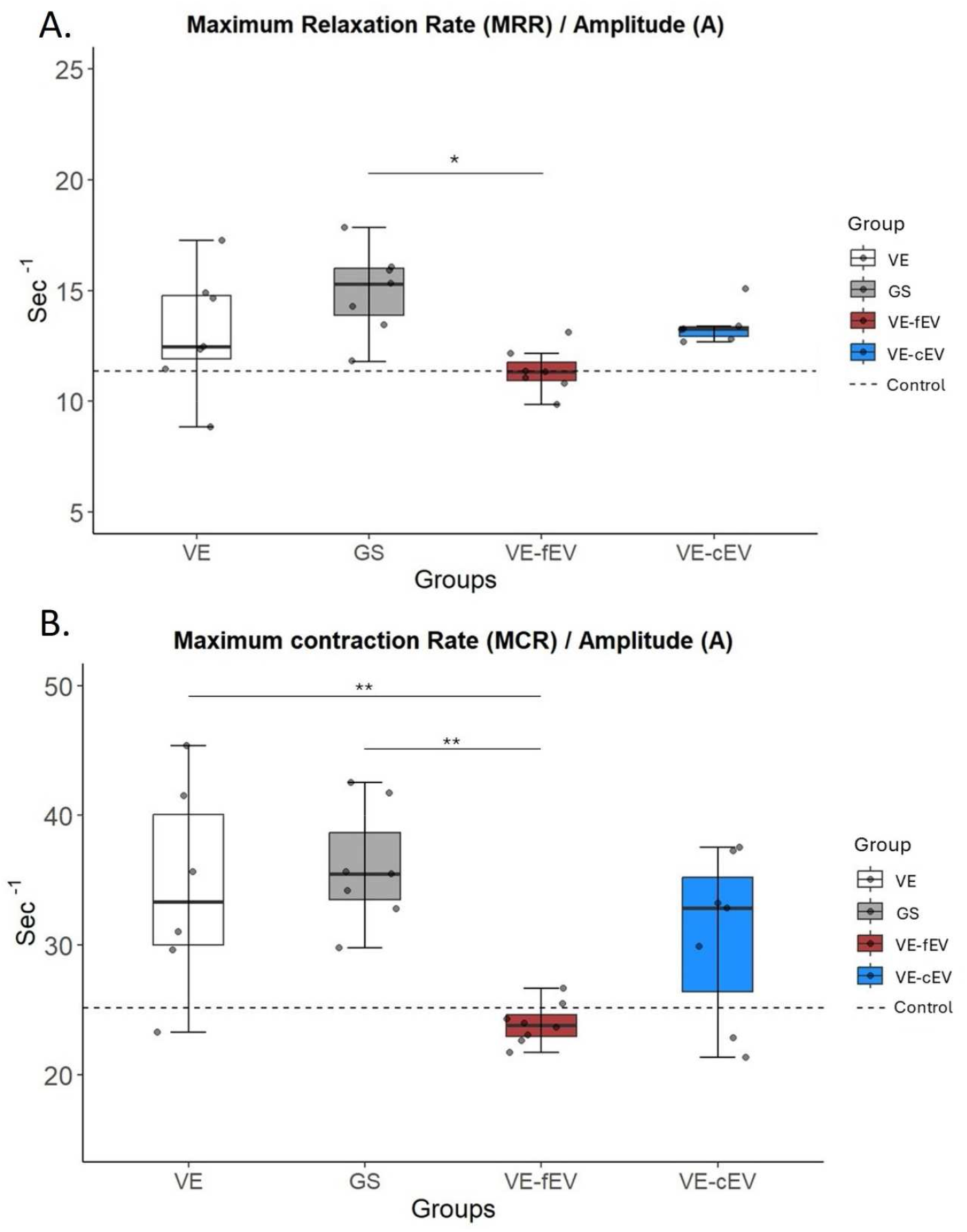
Muscle mechanical properties. **(A)** The fEV treated group displays a significant decrease in MCR/A ratio compared with VE and GS groups (p<0.01). Injection of fEVs allowed to maintain a phenotype close to Control group (dotted line). No difference was observed between VE-cEV group and VE, GS and VE-fEV groups. (B) Concerning the MRR/A ratio, only the VE-fEV treated group exhibits a significant reduction of the ratio when compared with the GS group (p<0.05) but no difference was observed with the two other groups (VE and VE-cEV). 1 symbol: p<0.05; 2 symbols: p<0.01; 3 symbols: p<0.001. Dotted line, mean value of Control; *GS, Gold Standard; VE, Vein; VE-cEV, Vein-cryoconserved extracellular vesicles; VE-fEV, Vein-freshly purified extracellular vesicles*.

### 3.3. OEMSC-derived EVs do not enhance nerve afferent response

#### Response to electrically induced fatigue

Afferent responses are similar in all vein-bridge animals, even when OEMSC-derived EVs (either freshly purified or cryoconserved) are grafted in the vein chamber (**Figure 3A**). No difference between the four groups is observed.

**Figure 3.**
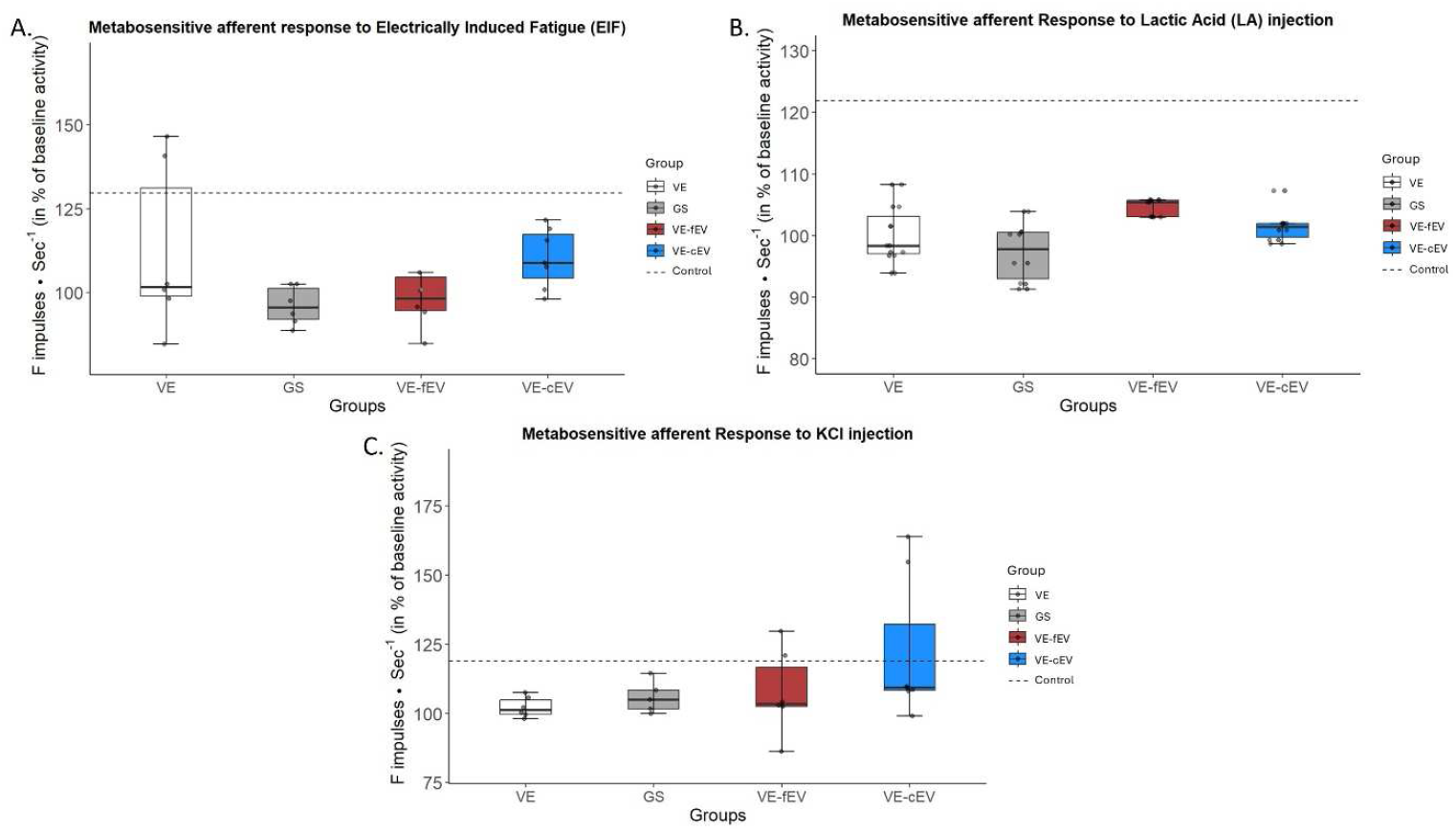
Nerve response to afferent stimulation. **(A)** Compared with GS group, responses associated to electrically induced fatigue were similar in VE animals, even when fEVs or cEVs were injected in the vein conduit. **(B)** No significant increase was observed in VE animals, even when either fEVs or cEVs were added in the vein chamber when compared with GS group, after injection of lactic acid. **(C)** The metabosensitive afferent responses to potassium chloride (KCl) were similar in all groups. OEMSC-derived EVs do not improve the recovery. 1 symbol: p<0.05; 2 symbols: p<0.01; 3 symbols: p<0.001. Dotted line, mean value of Control; *GS, Gold Standard; VE, Vein; VE-cEV, Vein-cryoconserved extracellular vesicles; VE-fEV, Vein-freshly purified extracellular vesicles*.

#### Response to lactic acid (LA) injection

No significant response is observed between experimental groups after injection of lactic acid (**Figure 3B**). Interestingly, when considering values in control rats (121.87 ± 7.315), no significant difference is observed between control and VE-fEV or VE-cEV groups, whereas a significant difference is observed between control and VE groups (p<0.05) or GS (p<0.05).

#### Response to Potassium chloride (KCl) injection

No significant response is observed in the VE, VE-fEV and VE-cEV rats compared with GS animals, after injection of KCl. Injected EVs do not improve recovery (**Figure 3C**).

### 3.4. OEMSC-derived EVs increase the number of axons

When the peroneal nerve defect is vein-bridged and filled with EVs, the number of axons significantly increase in the medial (p<0.001, for fEV; p<0.01, for cEV) and the distal parts (p<0.001, for fEV; p<0.01, for cEV) of the zone of injury, when compared with the vein conduit group (Figure 4A). Interestingly, the injection of freshly purified OEMSC-derived EVs (VE-fEV) into the vein leads to a significant increase in the number of axons in the medial part (p<0.01) when compared with the GS group (**Figure 4A**).

**Figure 4.**
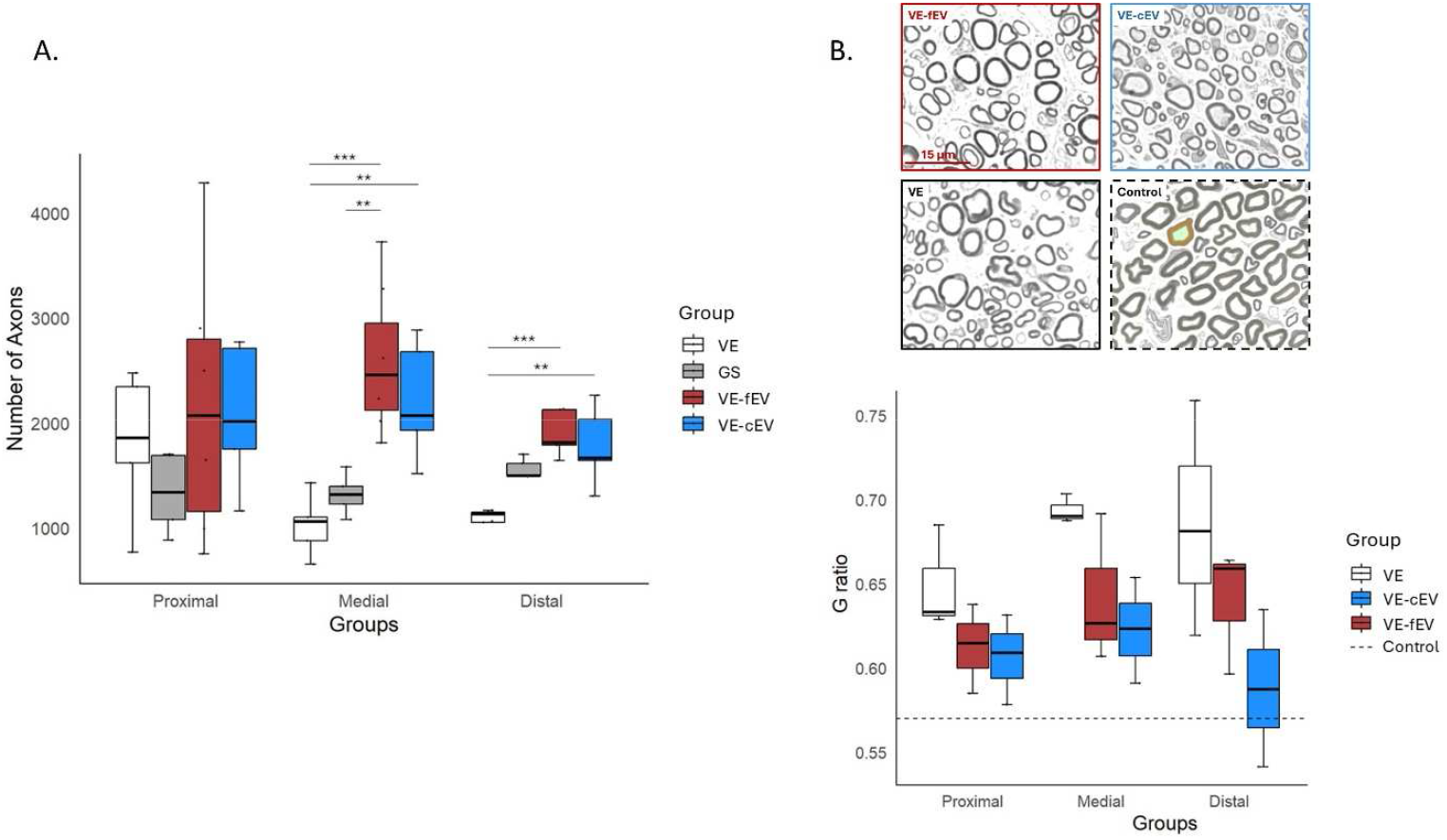
Histological analysis of the peroneal nerve. **(A)** Number of axons at the proximal, medial (vein bridge) and distal parts in the four groups of rats. The number increased in both the VE-fEV and VE-cEV groups, in the medial and proximal parts, when compared with VE group. When compared with GS, the number of axons only increased in the medial part for the fEV-treated animals. **(B)** Quantitative analysis of the G-ratio in the three groups observed. Above, representative TEM images of peroneal nerve in each group. Scale bar, 15 μm. No significant difference was observed between VE, VE-fEV and VE-cEV groups, even if the G-ratio decrease towards Control value (dotted line) for both fEV and cEV groups, in proximal, medial and distal parts of the damaged nerve. 1 symbol: p<0.05; 2 symbols: p<0.01; 3 symbols: p<0.001. Dotted line, mean value of Control; *GS, Gold Standard; VE, Vein; VE-cEV, Vein-cryoconserved extracellular vesicles; VE-fEV, Vein-freshly purified extracellular vesicles*.

### 3.4. OEMSC-derived EVs tend to improve myelination

The ratio of the inner axonal diameter to the total outer diameter (including the myelinated fiber) defines the g-ratio. The g-ratio is closely related to the myelin sheath around individual axons. **Figure 4B** shows a statistically non-significant trend towards an improved g-ratio for the animals transplanted with extracellular vesicles. However, no difference is observed between the three groups VE, VE-fEV and VE-cEV.

## 4. Discussion

The present study demonstrates that OEMSC-derived EVs induce axon sprouting, in distal and medial areas, similarly to the cells from which they are collected, compared with the Gold Standard group (reference in surgical practice). Interestingly, although freshly purified EVs (fEVs) induce a better regeneration than cryoconserved EVs, the latter, stored for 6 months at −80°C, still have a beneficial effect on nerve repair, close to those observed with the former. Injection of fEVs into a venous bridge connecting injured nerve endings promotes nerve regeneration and functional recovery more effectively in rat peroneal nerve injury model more effectively than Gold Standard treatment and the cEVs. These findings demonstrate that extracellular vesicles may be a novel agent for culturing and enhancing OEMSCs in neural repair after a peripheral nerve injury.

### 4.1. OE-MSC derived EVs improve motor functions in comparison with the Gold Standard treatment

#### Improvement of locomotion

Many studies demonstrate that EVs are the main regulators of the paracrine mechanism that mediates tissue regeneration ^38^. They carry proteins, lipids, and RNAs to recipient cells. The regenerative efficacy of MSC-derived EVs (adipose-derived MSCs, bone marrow MSCs, gingival MSC) was observed in various studies ^39^. The local application of rat bone marrow mesenchymal stem cell (BMSCs)-derived EVs in a rat model of sciatic nerve crush injury promotes nerve regeneration and functional recovery ^40^. The sciatic function index used to analyze the gait of injured rats confirms the improvement of the motor function in rats treated with BMSCs-derived EVs ^41^, 12 weeks after surgery. Similarly, adipocyte mesenchymal stem cell (asMSC)-derived EVs induce a faster functional recovery of the injured sciatic nerve ^42^ while human umbilical cord MSC-derived EVs (hUCMSC-EVs) increase the sciatic functional index, 8 weeks post-grafting ^43^.

Here, we demonstrate that both freshly purified and cryoconserved EVs allow full recovery of the locomotor function, at 9 and 12 weeks, respectively. Interestingly, OEMSC-derived EVs trigger significantly higher locomotor recovery than that observed in the Gold Standard (GS) group. In the case of lesions with loss of substance, the Gold Standard (nerve autograft) is considered by surgeons to be the reference treatment because it generates better results than end-to-end repairs carried out under tension, which lead to nerve ischemia ^44, 45^.

#### Improvement of muscle mechanical properties

Evoked muscle twitch provides information on the contractile properties as well as the quality of the regenerated efferent nerve fibers. It is widely recognized that when the peroneal nerve is damaged, the *Tibialis anterior* muscle, which is classified as a “fast” muscle, atrophies and transforms into a “slow” muscle ^46^. This change occurs when a vein alone is implanted and also in the Gold Standard care. Although the vein and the nerve autograft provide a supportive environment, neither of them is sufficient for the muscle to regain its original characteristics. Interestingly, injection of fEVs and cEVs enhances the mechanical properties of the muscle and allows it to bring back its original contractile phenotypes. In addition, freshly purified EVs completely restore the initial contractile properties of the muscle when compared with the Gold Standard.

#### Muscle mass is not correlated to function

Similar to the results obtained with OEMSC engraftment ^15^, recovery of locomotion is not associated with recovery of muscle mass. Studies show that hucMSCs and adMSC-derived exosomes lessen muscle atrophy following sciatic nerve injury, but EVs are directly injected in the *Gastrocnemius* muscle ^47, 48^, suggesting that additional injection of EVs in the *Tibialis anterior* muscle could improve and attenuate muscle atrophy. Moreover, the current study only considers the muscle weight, that is not a reflection of muscle recovery, since the latter is also influenced by other factors, such as the amount of intramuscular connective tissue ^49^ or phenotype of myosin chains ^50^. Further experiments should analyze the histology and morphology of the muscle fibers.

In conclusion, we can speculate that at the time of analysis (three months after surgery), the regenerated nerve fibers that have reached the target muscle may not be mature enough to restore muscle mass or its original characteristics. This could explain the miscorrelated results obtained for muscle atrophy and motor properties. To enhance the regeneration process, the use of electrical stimulation could be considered, as this technique has shown success in animal models and in humans ^51^.

### 4.2. OEMSC-derived EVs have same effects on sensitive fibers than Gold Standard

We show here that metabosensitive afferent responses are not improved for the four groups tested. The difference in recovery between the motor and sensory sides of the peroneal nerve may be a reflection of the duration of our study. Although it was shown that a 6 week post-surgical delay is sufficient to achieve functional motor recovery and histological outcomes in rats ^52^, metabosensitive connections might require more than 12 weeks ^53-55^.

### 4.3. OEMSC-derived EVs enhance axogenesis but not myelination

Improved walking behaviors are usually associated with increased axonal growth. In the present study, an obvious EV-related axon elongation/sprouting is at play. The present study demonstrates the positive add-on of EVs on axon regeneration. The number of axons is higher when EVs are injected regardless of storage. Both fEVs and cEVs increase axogenesis in the medial and distal parts. When compared with Gold Standard, only fEVs increase the number of axons in the medial part of the lesion. The g ratio observed in our study does not significantly differ from those obtained when an unfilled venous bridge is transplanted. Nevertheless, injection of fEVs or cEVs triggers the decrease of the g-ratio toward a ratio of 0.6 which is the mean value in peripheral axons ^56^. This trend suggests that injured axons are engaged in a remyelination process.

The purpose of the current study, considered as a pioneer work in the field of OEMSC-derived EVs, was not to unveil the underlying molecular mechanisms. However, previous studies using MSC-derived EVs allow to get a first picture of the involved molecules. It is established that EVs promote peripheral nerve recovery post-injury through various mechanisms, such as cell proliferation ^57^, migration ^58^, myelination ^59, 60^, and neurotrophic factor secretion ^61-63^. They carry soluble immunomodulatory mediators, including interleukin10 (IL-10) and Tumor Necrosis Factor (TNFα) that modulate neuroinflammation after a nerve trauma ^38, 62, 64^. Of interest, MSC-derived EVs are considered to be major immune mediators with more than 200 immunomodulatory proteins ^27^. In addition, MSC-EVs exert a pro-angiogenic effect *via* proteins associated with angiogenesis such as vascular endothelial growth factor (VEGF) and platelet-derived growth factor-D (PDGF-D), which promote angiogenesis ^40, 65^.

Other molecular mechanisms are supported and dependent on specific miRNA carried by MSC-derived EVs. For example, MSC-EVs transfer miRNA (miR-17-92 cluster, miR-221, miR-199B, miR-1489, miR-1350, miR-26a, miR133b, miR-26b and miR-22-3p) known to increase axonal growth and regeneration by activating PI3K/AKT, PTEN/mTOR or KPNA2 signaling pathways ^62, 63, 66-69^. They transfer angiogenic miRNA to enhance vascular regeneration by regulating AKT/eNOS, PI3K/AKT, TLR4/NF-KB or STAT3/AKT signaling pathways. BMSC-derived EVs transfer miR-1260a, −21-5p and −29b-3p to endothelial cells to promote vascular regeneration ^70-73^ as well as miR-210, −126, −125a, −423-5p, −31, −486, −26a- 5p and 147-3p ^63^. Moreover, miRNA-let7b, carried by hUMSC-EVS, activate TLR4/NF-KB/STAT3/AKT and thereby convert M1 macrophages into M2, leading to a downregulation of the inflammatory cytokines ^74, 75^.

### 4.4. Limitations and clinical perspectives

One obstacle for the use of EVs being their stability and storage, we compared the effect of freshly purified EVs *versus* cryoconserved EVs, which is recognized as the most effective storage method. Cryopreservation is routinely performed at the following temperatures: 4°C, − 80°C, and −196°C ^24^. However, storing exosomes at 4°C can negatively affect their biological activity and diminish their protein content. In contrast, –80°C is regarded as the ideal temperature, with the least detrimental effect on the morphology and composition of exosomes ^76^. In the current study, cEVs were cryopreserved at −80°C for 6 months. Now, it would be of interest to 1) establish a standardized process of preservation to gradually decrease the temperature at −80°C and 2) reduce the time of storage below 6 months to preserve the integrity of the molecules included in EVs.

Most pre-clinical *in vivo* studies on peripheral nerve regeneration assess the animal physiology during a period that does not exceed 3 months. However, it would be valuable to extend the post-trauma observation period to 6 and even 12 months. We observed a high number of axons in the medial and distal areas of the regenerating nerve. This result, in line with earlier findings in a rat nerve repair model, suggests that the number of distal axons may significantly increase during the first month and potentially doubles during the whole three-month period. However, axons that do not reach their target muscle tend to atrophy over time, with near-normal levels observed after two years ^77^. Similarly, it is plausible that, over the mid- or long-term, the *Tibialis anterior* muscle may regain its original mass and characteristics as nerve fibers mature. In additions, it could be planned to enhance and accelerate regeneration using additional strategies such as electrical stimulation or treadmill running. It has been shown that electrical stimulation promotes the reinnervation of motor and sensory motor and sensory neurons, enabling faster recovery ^78-80^ and is believed to have a beneficial effect on muscle atrophy and function ^54^.

## Conclusion

Further studies are needed to identify the components of the OEMSC-derived EVs in promoting nerve regeneration (RNA sequencing, proteomics analysis) and understand the molecular mechanisms underlying the nerve repair. Finally, such approaches will provide new insights to understand the role of OEMSCs in supporting nerve regeneration and exploring their potential therapeutic application in nerve injury repair.

## Acknowledgments

The electron microscopy experiments were performed by Aicha AOUANE, on the PiCSL-FBI core facility (IBDM, AMU-Marseille), member of the France-BioImaging national research infrastructure (ANR-10-INBS-04) and member of the Marseille Imaging Institute, an Excellence Initiative of Aix Marseille University A*MIDEX, a French “Investissements d’Avenir” programme (AMX 19 IET 002). The authors declare that they have not use AI-generated work in this manuscript.

## Authors contribution

MB and MS performed the experiments and generated figures and contributed in Writing Original Draft. MB and MS contributed equally to this work. MW and CJ provided guidance on the surgery methodology and in conceptualization. PM prepared nerve slices for immunohistochemistry. TM and PD contributed to electrophysiological experiments. FF contributed in conceptualization, design of the study, methodology and resources. GGC conceived and designed the study, cultured the OEMSC, purified the extracellular vesicles, guided the concept and data analysis and wrote the original draft. All the authors wrote and revised the manuscript.

## Statements and Declarations

### Ethical considerations

Experiments were performed according to the French law (Decrees and orders N°2013-118 of 01 February 2013, JORF n°0032) on animal care guidelines and after approval by animal Care Committees of Aix-Marseille Université (AMU) and Centre National de la Recherche Scientifique (CNRS). The authorization number granted by the French Ministry of Higher Education, Research, and Innovation (MESRI) is APAFIS#41012-2023021620519181 v5, entitled “*Réparation du nerf péronier lésé avec perte de substance à l’aide d’une greffe de vésicules extracellulaires issues des cellules souches olfactives humaines, chez le rat*” in August 2023. All persons were licensed to conduct live animal experiments and all room have a national authorization to accommodate animals (License n°B13.013.06). Furthermore, experiments were performed in accordance with the recommendations provided in the Guide for Care and Use of Laboratory Animals (U.S. Department of Health and Human Services, National Institutes of Health), with the directives 86/609/EEC and 010/63/EU of the European Parliament and of the Council of 24 November 1986 and of 22 September 2010, respectively, and with the ARRIVE (Animal Research: Reporting of *In Vivo* Experiments) guidelines.

### Consent to participate

The OEMSC were collected, previously in studies involving human participants. They were reviewed and approved by Comité de Protection des Personnes, file 2018-A00796-49. The patients/participants provided their written informed consent to participate in this study. Initiated in December 2018, a prospective study entitled NOSE (ClinicalTrials.gov Identifier: NCT04020367), aimed to validate the manufacture of olfactory stem cells, using good manufacturing practices, in order to use them as an Advanced Therapeutic Medicinal Product (ATMP) for repairing peripheral nerves.

### Consent for publication

Not applicable.

### Data availability

The raw data supporting the conclusions of this article will be made available by the authors, without undue reservation.

## References

1. Noble J, Munro CA, Prasad VS, et al. Analysis of upper and lower extremity peripheral nerve injuries in a population of patients with multiple injuries. J Trauma 1998; 45: 116–122. DOI: 10.1097/00005373-199807000-00025.

2. Robinson LR. Traumatic injury to peripheral nerves. Muscle Nerve 2000; 23: 863–873. DOI: 10.1002/(sici)1097-4598(200006)23:6<863::aid-mus4>3.0.co;2-0.

3. Li R, Liu Z, Pan Y, et al. Peripheral nerve injuries treatment: a systematic review. Cell Biochem Biophys 2014; 68: 449–454. DOI: 10.1007/s12013-013-9742-1.

4. Zou X, Dong Y, Alhaskawi A, et al. Techniques and graft materials for repairing peripheral nerve defects. Front Neurol 2023; 14: 1307883. 20240122. DOI: 10.3389/fneur.2023.1307883.

5. Murrell W, Féron F, Wetzig A, et al. Multipotent stem cells from adult olfactory mucosa. Dev Dyn 2005; 233: 496–515. DOI: 10.1002/dvdy.20360.

6. Graziadei GA and Graziadei PP. Neurogenesis and neuron regeneration in the olfactory system of mammals. II. Degeneration and reconstitution of the olfactory sensory neurons after axotomy. J Neurocytol 1979; 8: 197–213. DOI: 10.1007/bf01175561.

7. Girard SD, Deveze A, Nivet E, et al. Isolating nasal olfactory stem cells from rodents or humans. J Vis Exp 2011 20110822. DOI: 10.3791/2762.

8. Murrell W, Wetzig A, Donnellan M, et al. Olfactory mucosa is a potential source for autologous stem cell therapy for Parkinson’s disease. Stem Cells 2008; 26: 2183–2192. 20080605. DOI: 10.1634/stemcells.2008-0074.

9. Pandit SR, Sullivan JM, Egger V, et al. Functional effects of adult human olfactory stem cells on early-onset sensorineural hearing loss. Stem Cells 2011; 29: 670–677. DOI: 10.1002/stem.609.

10. Nivet E, Vignes M, Girard SD, et al. Engraftment of human nasal olfactory stem cells restores neuroplasticity in mice with hippocampal lesions. Journal of Clinical Investigation 2011; 121: 2808–2820. DOI: 10.1172/jci44489.

11. Delorme B, Nivet E, Gaillard J, et al. The human nose harbors a niche of olfactory ectomesenchymal stem cells displaying neurogenic and osteogenic properties. Stem Cells Dev 2010; 19: 853–866. DOI: 10.1089/scd.2009.0267.

12. Di Trapani M, Bassi G, Ricciardi M, et al. Comparative study of immune regulatory properties of stem cells derived from different tissues. Stem Cells Dev 2013; 22: 2990–3002. 20130809. DOI: 10.1089/scd.2013.0204.

13. Girard SD, Virard I, Lacassagne E, et al. From Blood to Lesioned Brain: An In Vitro Study on Migration Mechanisms of Human Nasal Olfactory Stem Cells. Stem Cells Int 2017; 2017: 1478606. 20170618. DOI: 10.1155/2017/1478606.

14. Bense F, Montava M, Duclos C, et al. Syngeneic Transplantation of Rat Olfactory Stem Cells in a Vein Conduit Improves Facial Movements and Reduces Synkinesis after Facial Nerve Injury. Plast Reconstr Surg 2020; 146: 1295–1305. DOI: 10.1097/prs.0000000000007367.

15. Bonnet M, Guiraudie-Capraz G, Marqueste T, et al. Immediate or Delayed Transplantation of a Vein Conduit Filled with Nasal Olfactory Stem Cells Improves Locomotion and Axogenesis in Rats after a Peroneal Nerve Loss of Substance. Int J Mol Sci 2020; 21 20200411. DOI: 10.3390/ijms21082670.

16. Fu Y, Karbaat L, Wu L, et al. Trophic Effects of Mesenchymal Stem Cells in Tissue Regeneration. Tissue Eng Part B Rev 2017; 23: 515–528. DOI: 10.1089/ten.TEB.2016.0365.

17. Sarvar DP, Effatpanah H, Akbarzadehlaleh P, et al. Mesenchymal stromal cell-derived extracellular vesicles: novel approach in hematopoietic stem cell transplantation. Stem Cell Res Ther 2022; 13: 202. 20220516. DOI: 10.1186/s13287-022-02875-3.

18. Thery C. Exosomes: secreted vesicles and intercellular communications. F1000 Biol Rep 2011; 3: 15. 20110701. DOI: 10.3410/B3-15.

19. Huang CW, Huang WC, Qiu X, et al. The Differentiation Stage of Transplanted Stem Cells Modulates Nerve Regeneration. Sci Rep 2017; 7: 17401. 20171212. DOI: 10.1038/s41598-017-17043-4.

20. Kirian RD, Steinman D, Jewell CM, et al. Extracellular vesicles as carriers of mRNA: Opportunities and challenges in diagnosis and treatment. Theranostics 2024; 14: 2265–2289. 20240311. DOI: 10.7150/thno.93115.

21. Yavuz B, Mutlu EC, Ahmed Z, et al. Applications of Stem Cell-Derived Extracellular Vesicles in Nerve Regeneration. Int J Mol Sci 2024; 25 20240528. DOI: 10.3390/ijms25115863.

22. Jung H, Jung Y, Seo J, et al. Roles of extracellular vesicles from mesenchymal stem cells in regeneration. Mol Cells 2024; 47: 100151. 20241113. DOI: 10.1016/j.mocell.2024.100151.

23. Kumar MA, Baba SK, Sadida HQ, et al. Extracellular vesicles as tools and targets in therapy for diseases. Signal Transduct Target Ther 2024; 9: 27. 20240205. DOI: 10.1038/s41392-024-01735-1.

24. Jeyaram A and Jay SM. Preservation and Storage Stability of Extracellular Vesicles for Therapeutic Applications. Aaps j 2017; 20: 1. 20171127. DOI: 10.1208/s12248-017-0160-y.

25. Riazifar M, Pone EJ, Lötvall J, et al. Stem Cell Extracellular Vesicles: Extended Messages of Regeneration. Annu Rev Pharmacol Toxicol 2017; 57: 125–154. 20161028. DOI: 10.1146/annurev-pharmtox-061616-030146.

26. Rahimian S, Najafi H, Webber CA, et al. Advances in Exosome-Based Therapies for the Repair of Peripheral Nerve Injuries. Neurochem Res 2024; 49: 1905–1925. 20240528. DOI: 10.1007/s11064-024-04157-1.

27. Lai RC, Tan SS, Teh BJ, et al. Proteolytic Potential of the MSC Exosome Proteome: Implications for an Exosome-Mediated Delivery of Therapeutic Proteasome. Int J Proteomics 2012; 2012: 971907. 20120718. DOI: 10.1155/2012/971907.

28. Feron F, Perry C, Girard SD, et al. Isolation of adult stem cells from the human olfactory mucosa. Methods Mol Biol 2013; 1059: 107–114. DOI: 10.1007/978-1-62703-574-3_10.

29. Jaloux C, Bonnet M, Vogtensperger M, et al. Human nasal olfactory stem cells, purified as advanced therapy medicinal products, improve neuronal differentiation. Front Neurosci 2022; 16: 1042276. 20221117. DOI: 10.3389/fnins.2022.1042276.

30. Bain JR, Mackinnon SE and Hunter DA. Functional evaluation of complete sciatic, peroneal, and posterior tibial nerve lesions in the rat. Plast Reconstr Surg 1989; 83: 129–138. DOI: 10.1097/00006534-198901000-00024.

31. Kaiser T, Allen HM, Kwon O, et al. MyelTracer: A Semi-Automated Software for Myelin g-Ratio Quantification. eNeuro 2021; 8 20210721. DOI: 10.1523/ENEURO.0558-20.2021.

32. Shapiro SS and Wilk MB. An analysis of variance test for normality (complete samples). Biometrika 1965; 52: 591–611. DOI: 10.1093/biomet/52.3-4.591.

33. Kruskal WH and Wallis WA. Use of Ranks in One-Criterion Variance Analysis. Journal of the American Statistical Association 1952; 47: 583–621. DOI: 10.2307/2280779.

34. Dunn OJ. Multiple Comparisons Using Rank Sums. Technometrics 1964; 6: 241–252. DOI: 10.1080/00401706.1964.10490181.

35. Scheirer CJ, Ray WS and Hare N. The analysis of ranked data derived from completely randomized factorial designs. Biometrics 1976; 32: 429–434.

36. Sokal R and Rohlf F. Biometry : the principles and practice of statistics in biological research / Robert R. Sokal and F. James Rohlf. 2013.

37. Mangiafico S. Summary and Analysis of Extension Program Evaluation in R. 2016.

38. Dong R, Liu Y, Yang Y, et al. MSC-Derived Exosomes-Based Therapy for Peripheral Nerve Injury: A Novel Therapeutic Strategy. BioMed Research International 2019; 2019: 1–12. DOI: 10.1155/2019/6458237.

39. Dogny C, André-Lévigne D, Kalbermatten DF, et al. Therapeutic Potential and Challenges of Mesenchymal Stem Cell-Derived Exosomes for Peripheral Nerve Regeneration: A Systematic Review. Int J Mol Sci 2024; 25 20240612. DOI: 10.3390/ijms25126489.

40. Ma Y, Ge S, Zhang J, et al. Mesenchymal stem cell-derived extracellular vesicles promote nerve regeneration after sciatic nerve crush injury in rats. Int J Clin Exp Pathol 2017; 10: 10032–10039. 20170901.

41. Zhang W, Fang XX, Li QC, et al. Reduced graphene oxide-embedded nerve conduits loaded with bone marrow mesenchymal stem cell-derived extracellular vesicles promote peripheral nerve regeneration. Neural Regen Res 2023; 18: 200–206. DOI: 10.4103/1673-5374.343889.

42. Bucan V, Vaslaitis D, Peck CT, et al. Effect of Exosomes from Rat Adipose-Derived Mesenchymal Stem Cells on Neurite Outgrowth and Sciatic Nerve Regeneration After Crush Injury. Mol Neurobiol 2019; 56: 1812–1824. 20180621. DOI: 10.1007/s12035-018-1172-z.

43. Ma Y, Dong L, Zhou D, et al. Extracellular vesicles from human umbilical cord mesenchymal stem cells improve nerve regeneration after sciatic nerve transection in rats. J Cell Mol Med 2019; 23: 2822–2835. 20190217. DOI: 10.1111/jcmm.14190.

44. Miyamoto Y. Experimental study of results of nerve suture under tension vs. nerve grafting. Plast Reconstr Surg 1979; 64: 540–549. DOI: 10.1097/00006534-197910000-00017.

45. Terzis J, Faibisoff B and Williams B. The nerve gap: suture under tension vs. graft. Plast Reconstr Surg 1975; 56: 166–170.

46. Marqueste T, Decherchi P, Dousset E, et al. Effect of muscle electrostimulation on afferent activities from tibialis anterior muscle after nerve repair by self-anastomosis. Neuroscience 2002; 113: 257–271. DOI: 10.1016/s0306-4522(02)00187-2.

47. Schilling BK, Schusterman MA, 2nd, Kim DY, et al. Adipose-derived stem cells delay muscle atrophy after peripheral nerve injury in the rodent model. Muscle Nerve 2019; 59: 603–610. 20190220. DOI: 10.1002/mus.26432.

48. Chen J, Zhu Y, Gao H, et al. HucMSCs Delay Muscle Atrophy After Peripheral Nerve Injury Through Exosomes by Repressing Muscle-Specific Ubiquitin Ligases. Stem Cells 2024; 42: 460–474. DOI: 10.1093/stmcls/sxae017.

49. Bueno CRS, Pereira M, Aparecido IF-J, et al. Comparative study between standard and inside-out vein graft techniques on sciatic nerve repair of rats. Muscular and functional analysis. Acta Cir Bras 2017; 32: 287–296. DOI: 10.1590/s0102-865020170040000005.

50. Marqueste T, Decherchi P, Desplanches D, et al. Chronic electrostimulation after nerve repair by self-anastomosis: effects on the size, the mechanical, histochemical and biochemical muscle properties. Acta Neuropathol 2006; 111: 589–600. 20060307. DOI: 10.1007/s00401-006-0035-2.

51. Gordon T and de Zepetnek JET. Motor unit and muscle fiber type grouping after peripheral nerve injury in the rat. Exp Neurol 2016; 285: 24–40. 20160902. DOI: 10.1016/j.expneurol.2016.08.019.

52. Brogan DM, Dy CJ, Rioux-Forker D, et al. Influences of Repair Site Tension and Conduit Splinting on Peripheral Nerve Reconstruction. Hand (N Y) 2022; 17: 1048–1054. 20201224. DOI: 10.1177/1558944720974117.

53. Decherchi P, Vuillon-Cacciutolo G, Darques JL, et al. Changes in afferent activities from tibialis anterior muscle after nerve repair by self-anastomosis. Muscle Nerve 2001; 24: 59–68. DOI: 10.1002/1097-4598(200101)24:1<59::aid-mus7>3.0.co;2-s.

54. Marqueste T, Alliez JR, Alluin O, et al. Neuromuscular rehabilitation by treadmill running or electrical stimulation after peripheral nerve injury and repair. J Appl Physiol (1985) 2004; 96: 1988–1995. 20031121. DOI: 10.1152/japplphysiol.00775.2003.

55. Pertici V, Laurin J, Féron F, et al. Functional recovery after repair of peroneal nerve gap using different collagen conduits. Acta Neurochir (Wien) 2014; 156: 1029–1040. 20140205. DOI: 10.1007/s00701-014-2009-9.

56. Chomiak T and Hu B. What is the optimal value of the g-ratio for myelinated fibers in the rat CNS? A theoretical approach. PLoS One 2009; 4: e7754. 20091113. DOI: 10.1371/journal.pone.0007754.

57. Lopez-Verrilli MA, Caviedes A, Cabrera A, et al. Mesenchymal stem cell-derived exosomes from different sources selectively promote neuritic outgrowth. Neuroscience 2016; 320: 129–139. 20160203. DOI: 10.1016/j.neuroscience.2016.01.061.

58. Mao Q, Nguyen PD, Shanti RM, et al. Gingiva-Derived Mesenchymal Stem Cell-Extracellular Vesicles Activate Schwann Cell Repair Phenotype and Promote Nerve Regeneration. Tissue Eng Part A 2019; 25: 887–900. 20181228. DOI: 10.1089/ten.TEA.2018.0176.

59. Clark K, Zhang S, Barthe S, et al. Placental Mesenchymal Stem Cell-Derived Extracellular Vesicles Promote Myelin Regeneration in an Animal Model of Multiple Sclerosis. Cells 2019; 8 20191123. DOI: 10.3390/cells8121497.

60. Clark KC, Wang D, Kumar P, et al. The Molecular Mechanisms Through Which Placental Mesenchymal Stem Cell-Derived Extracellular Vesicles Promote Myelin Regeneration. Adv Biol (Weinh) 2022; 6: e2101099. 20220113. DOI: 10.1002/adbi.202101099.

61. Klimovich P, Rubina K, Sysoeva V, et al. New Frontiers in Peripheral Nerve Regeneration: Concerns and Remedies. Int J Mol Sci 2021; 22 20211213. DOI: 10.3390/ijms222413380.

62. Li Q, Zhang F, Fu X, et al. Therapeutic Potential of Mesenchymal Stem Cell-Derived Exosomes as Nanomedicine for Peripheral Nerve Injury. Int J Mol Sci 2024; 25 20240718. DOI: 10.3390/ijms25147882.

63. Yang S, Sun Y and Yan C. Recent advances in the use of extracellular vesicles from adipose-derived stem cells for regenerative medical therapeutics. J Nanobiotechnology 2024; 22: 316. 20240606. DOI: 10.1186/s12951-024-02603-4.

64. Pegtel DM and Gould SJ. Exosomes. Annu Rev Biochem 2019; 88: 487–514. DOI: 10.1146/annurev-biochem-013118-111902.

65. Du W, Zhang K, Zhang S, et al. Enhanced proangiogenic potential of mesenchymal stem cell-derived exosomes stimulated by a nitric oxide releasing polymer. Biomaterials 2017; 133: 70–81. 20170417. DOI: 10.1016/j.biomaterials.2017.04.030.

66. Li M, Lei H, Xu Y, et al. Exosomes derived from mesenchymal stem cells exert therapeutic effect in a rat model of cavernous nerves injury. Andrology 2018; 6: 927–935. 20180716. DOI: 10.1111/andr.12519.

67. Namini MS, Daneshimehr F, Beheshtizadeh N, et al. Cell-free therapy based on extracellular vesicles: a promising therapeutic strategy for peripheral nerve injury. Stem Cell Res Ther 2023; 14: 254. 20230919. DOI: 10.1186/s13287-023-03467-5.

68. Xin H, Katakowski M, Wang F, et al. MicroRNA cluster miR-17-92 Cluster in Exosomes Enhance Neuroplasticity and Functional Recovery After Stroke in Rats. Stroke 2017; 48: 747–753. DOI: 10.1161/strokeaha.116.015204.

69. Chen Y, Tian Z, He L, et al. Exosomes derived from miR-26a-modified MSCs promote axonal regeneration via the PTEN/AKT/mTOR pathway following spinal cord injury. Stem Cell Res Ther 2021; 12: 224. 20210405. DOI: 10.1186/s13287-021-02282-0.

70. Zhang L, Ouyang P, He G, et al. Exosomes from microRNA-126 overexpressing mesenchymal stem cells promote angiogenesis by targeting the PIK3R2-mediated PI3K/Akt signalling pathway. J Cell Mol Med 2021; 25: 2148–2162. 20201221. DOI: 10.1111/jcmm.16192.

71. Hu H, Hu X, Li L, et al. Exosomes Derived from Bone Marrow Mesenchymal Stem Cells Promote Angiogenesis in Ischemic Stroke Mice via Upregulation of MiR-21-5p. Biomolecules 2022; 12 20220624. DOI: 10.3390/biom12070883.

72. Zheng J, Zhang X, Cai W, et al. Bone Marrow Mesenchymal Stem Cell-Derived Exosomal microRNA-29b-3p Promotes Angiogenesis and Ventricular Remodeling in Rats with Myocardial Infarction by Targeting ADAMTS16. Cardiovasc Toxicol 2022; 22: 689–700. 20220614. DOI: 10.1007/s12012-022-09745-7.

73. Yu M, Liu W, Li J, et al. Exosomes derived from atorvastatin-pretreated MSC accelerate diabetic wound repair by enhancing angiogenesis via AKT/eNOS pathway. Stem Cell Res Ther 2020; 11: 350. 20200812. DOI: 10.1186/s13287-020-01824-2.

74. Ti D, Hao H, Tong C, et al. LPS-preconditioned mesenchymal stromal cells modify macrophage polarization for resolution of chronic inflammation via exosome-shuttled let-7b. J Transl Med 2015; 13: 308. 20150919. DOI: 10.1186/s12967-015-0642-6.

75. Sun G, Li G, Li D, et al. hucMSC derived exosomes promote functional recovery in spinal cord injury mice via attenuating inflammation. Mater Sci Eng C Mater Biol Appl 2018; 89: 194–204. 20180410. DOI: 10.1016/j.msec.2018.04.006.

76. Aldali F, Deng C, Nie M, et al. Advances in therapies using mesenchymal stem cells and their exosomes for treatment of peripheral nerve injury: state of the art and future perspectives. Neural Regen Res 2024 20241022. DOI: 10.4103/nrr.Nrr-d-24-00235.

77. Mackinnon SE, Dellon AL and O’Brien JP. Changes in nerve fiber numbers distal to a nerve repair in the rat sciatic nerve model. Muscle Nerve 1991; 14: 1116–1122. DOI: 10.1002/mus.880141113.

78. Al-Majed AA, Neumann CM, Brushart TM, et al. Brief electrical stimulation promotes the speed and accuracy of motor axonal regeneration. J Neurosci 2000; 20: 2602–2608. DOI: 10.1523/JNEUROSCI.20-07-02602.2000.

79. Brushart TM, Jari R, Verge V, et al. Electrical stimulation restores the specificity of sensory axon regeneration. Exp Neurol 2005; 194: 221–229. DOI: 10.1016/j.expneurol.2005.02.007.

80. Gordon T and Borschel GH. The use of the rat as a model for studying peripheral nerve regeneration and sprouting after complete and partial nerve injuries. Exp Neurol 2017; 287: 331–347. 20160118. DOI: 10.1016/j.expneurol.2016.01.014.

